# Ancestral absence of electron transport chains in Patescibacteria and DPANN

**DOI:** 10.1101/2020.04.07.029462

**Authors:** Jacob P. Beam, Eric D. Becraft, Julia M. Brown, Frederik Schulz, Jessica K. Jarett, Oliver Bezuidt, Nicole J. Poulton, Kayla Clark, Peter F. Dunfield, Nikolai V. Ravin, John R. Spear, Brian P. Hedlund, Konstantinos A. Kormas, Stefan M. Sievert, Mostafa S. Elshahed, Hazel A. Barton, Matthew B. Stott, Jonathan A. Eisen, Duane P. Moser, Tullis C. Onstott, Tanja Woyke, Ramunas Stepanauskas

## Abstract

Recent discoveries suggest that the candidate superphyla Patescibacteria and DPANN constitute a large fraction of the phylogenetic diversity of Bacteria and Archaea. Their small genomes and limited coding potential have been hypothesized to be ancestral adaptations to obligate symbiotic lifestyles. To test this hypothesis, we performed cell-cell association, genomic, and phylogenetic analyses on 4,829 individual cells of Bacteria and Archaea from 46 globally distributed surface and subsurface field samples. This confirmed the ubiquity and abundance of Patescibacteria and DPANN in subsurface environments, the small size of their genomes and cells, and the divergence of their gene content from other Bacteria and Archaea. Our analyses suggest that most Patescibacteria and DPANN in the studied subsurface environments do not form specific physical associations with other microorganisms. These data also suggest that their unusual genomic features and prevalent auxotrophies may be a result of minimal cellular energy transduction mechanisms that potentially precede the evolution of respiration, thus relying solely on fermentation for energy conservation.

## Introduction

Cultivation-independent research tools have revealed the coding potential of numerous, deep branches of Bacteria and Archaea that were unknown until recently (Wrighton et al., 2012; Rinke et al., 2013; Brown et al., 2015; Castelle et al., 2015) Among them, the candidate bacterial superphylum Patescibacteria (also known as Candidate Phyla Radiation, CPR) and archaeal superphylum DPANN have garnered particular attention, as they appear to constitute a large fraction of microbial diversity in subsurface and various other environments (Brown et al., 2015; Hug et al., 2016; Dombrowski et al., 2019). Patescibacteria and DPANN are characterized by small genomes and cell sizes, and predicted minimal biosynthetic and metabolic potential (Wrighton et al., 2012; Luef et al., 2015; Castelle and Banfield, 2018). They also appear to have slow metabolism, as indicated by low per-cell ribosome counts (Luef et al., 2015) and slow estimated genome replication rates (Brown et al., 2016). Host-dependent endo- and ecto-symbioses have been observed in several Patescibacteria (Gong et al., 2014; He et al., 2015; Cross et al., 2019) and the Nanoarchaeota and Nanohaloarchaeota phyla within DPANN (Huber et al., 2002; Podar et al., 2013; Munson-McGee et al., 2015; Jarett et al., 2018; Hamm et al., 2019). As a result, it has been posited that the unusual biological features of Patescibacteria and DPANN reflect ancestral adaptations to symbiotic lifestyles (Castelle et al., 2018; Dombrowski et al., 2019). However, direct evidence of symbiosis in Patescibacteria and DPANN is limited to a small number of narrow phylogenetic groups inhabiting surface environments and, in the case of Patescibacteria, dependent on eukaryotic hosts (Gong et al., 2014) or eukaryotic host systems (He et al., 2015; Cross et al., 2019) (i.e., mammalian oral cavities), which suggests relatively recent adaptations.

Here, we performed physical cell-cell association, genomic, and phylogenetic analyses on 4,829 of individual microbial cells from 46 globally distributed and environmentally diverse locations to gain additional insights into the unusual biological features of Patescibacteria and DPANN. Consistent with prior reports, we found these two superphyla abundant in many subsurface environments, and also confirm their consistently small cell and genome sizes. Our single cell genomic and biophysical observations do not support the prevailing view that Patescibacteria and DPANN are dominated by symbionts (Castelle et al., 2018; Dombrowski et al., 2019). Instead, based on the apparent lack of genes for complete electron transport systems, we hypothesize that these two superphyla have not evolved the capacity for respiration and therefore rely on fermentative metabolisms for energy conservation. Although complex metabolic interdependencies are a rule rather than exception in natural microbiomes (Zengler and Zaramela, 2018), the predicted fermentative energy conservation and limited biosynthetic potential (Castelle et al., 2018; Dombrowski et al., 2019) of Patescibacteria and DPANN may define a highly communal lifestyle of these two superphyla and provide explanation for the extreme difficulty in obtaining them in pure culture.

## Materials and Methods

### Field sample collection

Field samples were collected from a global set of diverse environments that were found to contain candidate phyla of Bacteria and Archaea in prior studies (Rinke et al., 2013; Thomas et al., 2013; Moser et al., 2015; Becraft et al., 2017; Hershey et al., 2018; Sackett, 2018; Sackett et al., 2018, 2019). Immediately after collection, samples were amended with sterile 5% glycerol and 1 mM EDTA (final concentrations) and stored at −80 °C. Field sample metadata is located with each individual SAG in Table S1.

### Single amplified genome (SAG) generation, sequencing, and *de novo* assembly

SAG generation and sequencing were performed by Bigelow Laboratory for Ocean Sciences Single Cell Genomics Center (SCGC) and U.S. Department of Energy Joint Genome Institute (JGI) (Table S1). At SCGC, field samples were stained with SYTO-9 nucleic acids stain (Thermo Fisher Scientific), separated using fluorescence-activated cell sorting (FACS), lysed using a combination of freeze-thaw and alkaline treatment, and their genomic DNA was amplified using WGA-X in a cleanroom, as previously described (Stepanauskas et al., 2017). For sorting of cells with active oxidoreductases, the Beatrix field sample (plate AG-274) was pre-incubated with the RedoxSensor Green stain (Thermo Fisher Scientific) following manufacturer’s instructions. During cell sorting, cell size estimates were performed using calibrated index FACS (Stepanauskas et al., 2017). All SAGs generated at SCGC were subject to Low Coverage Sequencing (LoCoS) using a modified Nextera library preparation protocol and NextSeq 500 (Illumina) sequencing instrumentation (Stepanauskas et al., 2017). This resulted in a variable number of 2×150bp reads per SAG, with an average of ∼300k. The reads were de novo assembled using a customized workflow utilizing SPAdes (Bankevich et al., 2012), as previously described (Stepanauskas et al., 2017). The quality of the sequencing reads was assessed using FastQC and the quality of the assembled genomes (contamination and completeness) was assessed using checkM (Parks et al., 2015) and tetramer frequency analysis (Woyke et al., 2009). This SAG generation, sequencing and assembly workflow was previously evaluated for assembly errors using three bacterial benchmark cultures with diverse genome complexity and GC content (%), indicating no non-target and undefined bases in the assemblies and average frequencies of mis-assemblies, indels and mismatches per 100 kbp being 1.5, 3.0 and 5.0 (Stepanauskas et al., 2017). Functional annotation was first performed using Prokka (Seemann, 2014) with default Swiss-Prot databases supplied by the software. Prokka was run a second time with a custom protein annotation database built from compiling Swiss-Prot (Bateman et al., 2017) entries for Archaea and Bacteria. The uniquely barcoded sequencing libraries of SAGs belonging to candidate divisions were combined, in equal proportions, into 48-library pools and shipped to JGI for deeper sequencing with NextSeq 500 (Illumina) in 2×150 bp mode. Quality filtering of raw reads was performed with BBTools v.37, read normalization with BBNorm, and error correction with Tadpole (http://bbtools.jgi.doe.gov). The resulting reads were assembled with SPAdes (Nurk et al., 2013) (v3.9.0, --phred-offset 33 –sc -k 22,55,95 –12), and 200 bp was trimmed from the ends of assembled contigs, after which contigs with read coverage < 2 or < 2 kbp in length were discarded. Assemblies were annotated according to IMG standard protocols (Huntemann et al., 2016; Chen et al., 2019). All SAGs are publicly available in IMG/M (Chen et al., 2019), and can be found under their GOLD analysis project identifiers in Table S1.

### Identification of heterogenous DNA sources

The 16S ribosomal RNA gene was identified in SAGs by searching them individually using cmsearch, which is part of the infernal package (Nawrocki and Eddy, 2013), using the bacterial 16S rRNA Rfam covariance model (rfam.xfam.org/family/RF00177). This method is particularly helpful in predicting 16S rRNA genes in Patescibacteria and DPANN, which can often have introns in their 16S rRNA genes (Brown, 2015). Taxonomic assignments to these 16S rRNA genes were conducted using “classify.seqs” within mothur (Schloss et al., 2009) version 1.41.3 against the Silva 132 reference database and taxonomy file (Quast et al., 2013). The resulting taxonomy file was used to search for SAGs that contained two 16S rRNA genes that had different taxonomic phylum-level assignments and were marked as putative co-sorts; those that did not have two 16S rRNA genes were marked as single sorts. The checkM (Parks et al., 2015) contamination estimates were used to determine SAGs that had high values of potential genome admixture (e.g., two different cellular origins). A Chi-squared test was performed in R using the “chisq.test” function on potential co-sorted and single sorted SAGs, and Pearson’s residuals were retrieved from the output of this test and used to calculate the percent contribution to each X^2^ statistic, and plotted using the “corrplot” package in R Studio.

### Genomes from prior studies

A total of 1,025 publicly available SAGs, metagenome bins, and isolate genomes (Table S2) were used in this study from the Integrated Microbial Genomes and Microbiomes (IMG/M) database (Chen et al., 2019) (genomes accessed April 2018). These genomes were selected by clustering the RNA polymerase COG0086 protein sequence at 70% identity, and if there were similar genomes at the 70% identity threshold, the one with the most complete set of 56 single copy proteins was chosen as representative. Phylum-level classification and symbiotic lifestyle assignments were exported from IMG/M. In cases were IMG/M lacked lifestyle assignments, manual literature searches of organism names were used to determine whether they have documented symbiotic relationships.

### Concatenated single copy protein phylogeny

A set of 56 universal single copy marker proteins (Eloe-Fadrosh et al., 2016; Yu et al., 2017) was used to build a phylogenetic tree for the newly generated SAGs and MAGs and a representative set of bacteria and archaea based on publicly available microbial genomes in IMG/M (Chen et al., 2019) (genomes accessed in April 2018). Marker proteins were identified with hmmsearch (Eddy, 2011) version 3.1b2, using a specific Hidden Markov Model for each of the markers. Genomes for which 5 or more different marker proteins could be identified were included in the tree. For every marker protein, alignments were built with MAFFT (Nakamura et al., 2018) v7.294b and subsequently trimmed with BMGE (Criscuolo and Gribaldo, 2010) v1.12 using BLOSUM30. Single protein alignments were concatenated and maximum likelihood phylogenies inferred with FastTree2 (Price et al., 2010) using the options: -spr 4 -mlacc 2 - slownni -lg (for archaea) and -spr 4 -mlacc 2 -slownni -lg (for bacteria).

### Clusters of orthologous groups principal components analysis

Clusters of orthologous groups (COGs) were assigned to SAG (Table S1) and reference genome (Table S2) predicted protein sequences using reverse position-specific blast (rpsblast) (Altschul et al., 1997) with an e-value cutoff of 1e-5 and the cdd2cog script (https://github.com/aleimba/bac-genomics-scripts/tree/master/cdd2cog). Genomes that were used for the principal component analysis (PCA) had completeness estimates greater than or equal to 30%, and contained 16S rRNA genes for unambiguous phylum-level classification. Eigenvector values were calculated in RStudio (RStudio Team, 2016) version 1.1.463 using the cmdscale function from relative abundances of the different COG categories expressed as a percent out of the total number of assigned COGs. PCA plots were visualized with ggplot2 (Wickham, 2016) in RStudio (RStudio Team, 2016). A Wilcoxon test was performed in RStudio using the “wilcox.test” function to determine statistical differences between principal components among the different clusters discussed in the main text. The color scheme for these plots is based on the Color Universal Design (https://jfly.uni-koeln.de/color/), and should be distinguishable by all types of vision. This color scheme was used throughout all the figures in the manuscript.

### Coding sequence density

Coding sequences (CDS) for SAGs and reference genomes were predicted using Prodigal (Hyatt et al., 2010) version 2.6.3. The initial analysis of prokka CDS density revealed that numerous SAGs and reference genomes had very low coding densities. Prokka utilizes the code 11 translation table by default, and many of these genomes could potentially use stop codons in place of canonical codons (Wrighton et al., 2012; Rinke et al., 2013). We determined the correct translation table to utilize for each genome by comparing the total CDS length from Code 11 and Code 25 predictions, and if the Code 11 total CDS length was greater than the Code 25 total CDS length, then the total length from Code 11 was used in the coding density calculation. If the opposite was true, then the Code 25 total CDS length was used. The coding density was calculated by dividing the total CDS sequence by the total assembly size.

### Oxygen reductase identification

A published heme copper oxidase subunit I database (Sousa et al., 2011) from bacteria and archaea was used as a database with blastp (Altschul et al., 1990) with an e-value cutoff of 1e-10 using the SAG and reference genomes as queries. The original database file had to be de-replicated (i.e., removing 100% identical sequences) using the dedupe.sh script, which is part of the BBMap package (https://github.com/BioInfoTools/BBMap). The sole crystal structure sequence for the bd-ubiquinol oxidase subunit A from *Geobacillus thermodentrificans* (Safarian et al., 2016) was used as a database for a blastp (Altschul et al., 1990) search using the SAGs and reference genomes as queries with an e-value cutoff of 1e-10.

### Oxygen reductase horizontal gene transfer

The protein sequences identified from the above section were retrieved from SAGs using the grep function from the list of sequence file headers from the above analysis in the SeqKit package (Shen et al., 2016). Reference protein sequences for Patescibacteria were retrieved via the blastp server using the Patescibacteria SAG HCO sequences as queries and selecting for hits only from sequences that were assigned to Patescibacteria and/or Candidate Phyla Radiation. Other reference sequences for Patescibacteria were retrieved by manual literature searches from relevant studies (Nelson and Stegen, 2015; León-Zayas et al., 2017; Castelle et al., 2018). The search for Patescibacteria HCOs revealed that they only encoded for the low-affinity Type A HCO, and all subsequent phylogenetic analyses focused solely on this HCO type. The multi-fasta file containing all HCO sequences was filtered for sequences that were greater than 400 amino acids in length, and aligned with mafft (Nakamura et al., 2018) using the “--auto” option and the resulting alignment was trimmed with trimal (Capella-Gutiérrez et al., 2009) to remove gaps using the “-gappyout” option. A maximum likelihood phylogenetic tree was created using FastTree (Price et al., 2010) using the LG model of amino acid evolution. No DPANN genome to date has had a positive identification of an HCO subunit I. The methodology for the HCO phylogeny was repeated for the bd-ubiquinol oxygen reductases. Phylogenetic trees were visualized and annotated using the online Interactive Tree of Life tool (Letunic and Bork, 2019).

### Oxidoreductase annotation and abundance

Enzyme Commission 1 (EC1) class family proteins (i.e., oxidoreductases) were predicted from the SAGs and reference genomes using the prokka “genome.tsv” annotation files. The total number of predicted protein sequences annotated as EC1 for each genome was divided by the total number of predicted protein sequences to provide the percent of protein encoding genes that were predicted to be oxidoreductases. This allows for a direct comparison of all the genomes that exhibited wide ranges in completeness estimates.

### KEGG orthology assignment of electron transport chain proteins

The Kyoto Encyclopedia of Gene and Genomes (KEGG) orthology (KO) annotations were assigned using KofamKOALA (Aramaki et al., 2019), which uses hmmsearch (Eddy, 2011) against curated hidden Markov model (HMM) KO profiles. Only KO profiles related to energy transduction oxidoreductases were used to search the genomes in this study, which were extracted from Supplemental Table 1 in Jelen et al. (2016). Sequences were identified as positive hits if their score was greater than or equal to 50% of the sequence threshold value as calculated in KofamKOALA.

### 16S ribosomal RNA gene phylogeny

16S rRNA gene sequences predicted using cmsearch (Nawrocki and Eddy, 2013) were filtered for sequences that were greater than or equal to 1200 bp using bioawk (https://github.com/lh3/bioawk). Sequences that were 100% identical were removed using dedupe.sh (https://github.com/BioInfoTools/BBMap). Sequences were then aligned using ssu-align (Nawrocki, 2009), which produces two separate alignment files for Bacteria and Archaea. Next, ambiguously aligned positions were removed using ssu-mask, and sequences were re-checked to ensure that the masked alignment contained sequences that were greater than or equal to 1200 bp. Sequences that did not meet these threshold requirements were removed from the alignment file using ssu-mask with the “--seq-r” option and list of sequences to remove. The Stockholm alignment file was converted to an aligned fasta file using ssu-mask with the “--stk2afa” option. The masked and filtered alignment files for Bacteria and Archaea were used to create phylogenetic trees using maximum likelihood reconstruction with FastTree (Price et al., 2010) with the following parameters: “-nt -gtr -cat 20 -gamma”. Both trees were visualized and annotated using the Interactive Tree of Life (Letunic and Bork, 2019).

## Results and Discussion

### Global presence of Patescibacteria and DPANN in subsurface environments

To improve our understanding of the deep genealogy of Bacteria and Archaea, we sequenced 4,829 single amplified genomes (SAGs; Table S1) of previously under-sampled microbial lineages from 46 globally distributed field sites (Figure 1; Table S1). These sites were chosen based on 16S rRNA gene amplicon screens that were enriched in bacterial and archaeal candidate phyla. A maximum likelihood phylogenetic tree of concatenated single-copy proteins (SCP) (Figure 2) positioned 22% and 4% of SAGs within Patescibacteria (n=492) and DPANN archaea (n=81). The concatenated SCP phylogenetic tree revealed the separation of Patescibacteria and DPANN from other Bacteria and Archaea, respectively, which corroborates other phylogenetic reconstructions using diverse sets of single copy proteins and phylogenetic tools (Rinke et al., 2013; Brown et al., 2015; Hug et al., 2016; Williams et al., 2017; Castelle et al., 2018; Dombrowski et al., 2019). Patescibacteria comprised a median relative abundance of 13% (range=0-81%) and DPANN comprised a median abundance of 7.5% (range=0-23%) in 33 analyzed environmental sites, with elevated abundances in deep-sourced aquifer environments (Figure 3). Most of the Patescibacteria and DPANN SAGs originated from 13 continental subsurface sites in Africa, Asia, and North America (Table S1). These results confirm that Patescibacteria and DPANN are globally abundant members of subsurface microbial communities, expanding on the prior genomic studies that were predominantly based on a small number of study locations in North America (Rinke et al., 2013; Luef et al., 2015; Castelle et al., 2018).

**Figure 1.**
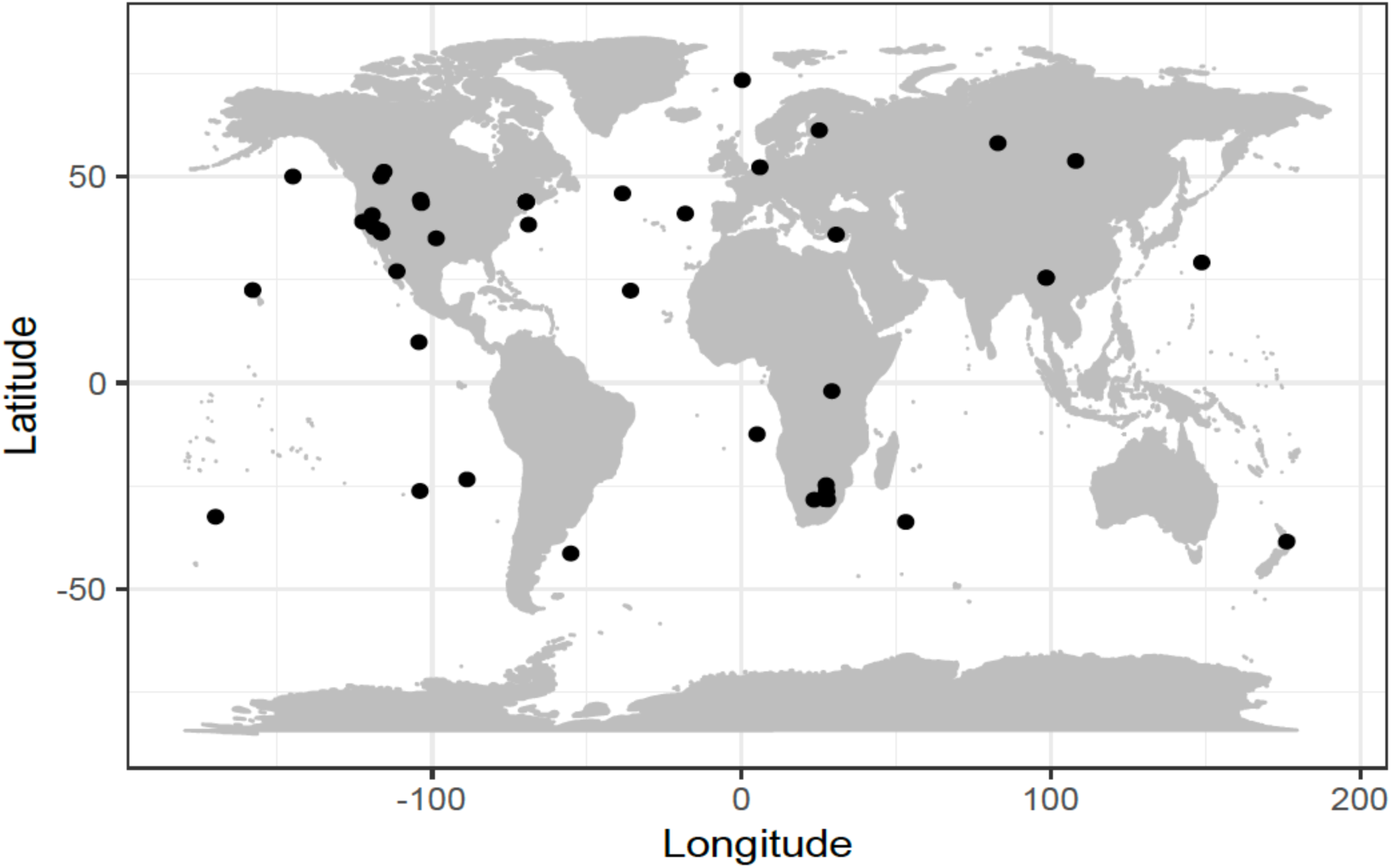
Geographic locations of sample collection sites.

**Figure 2.**
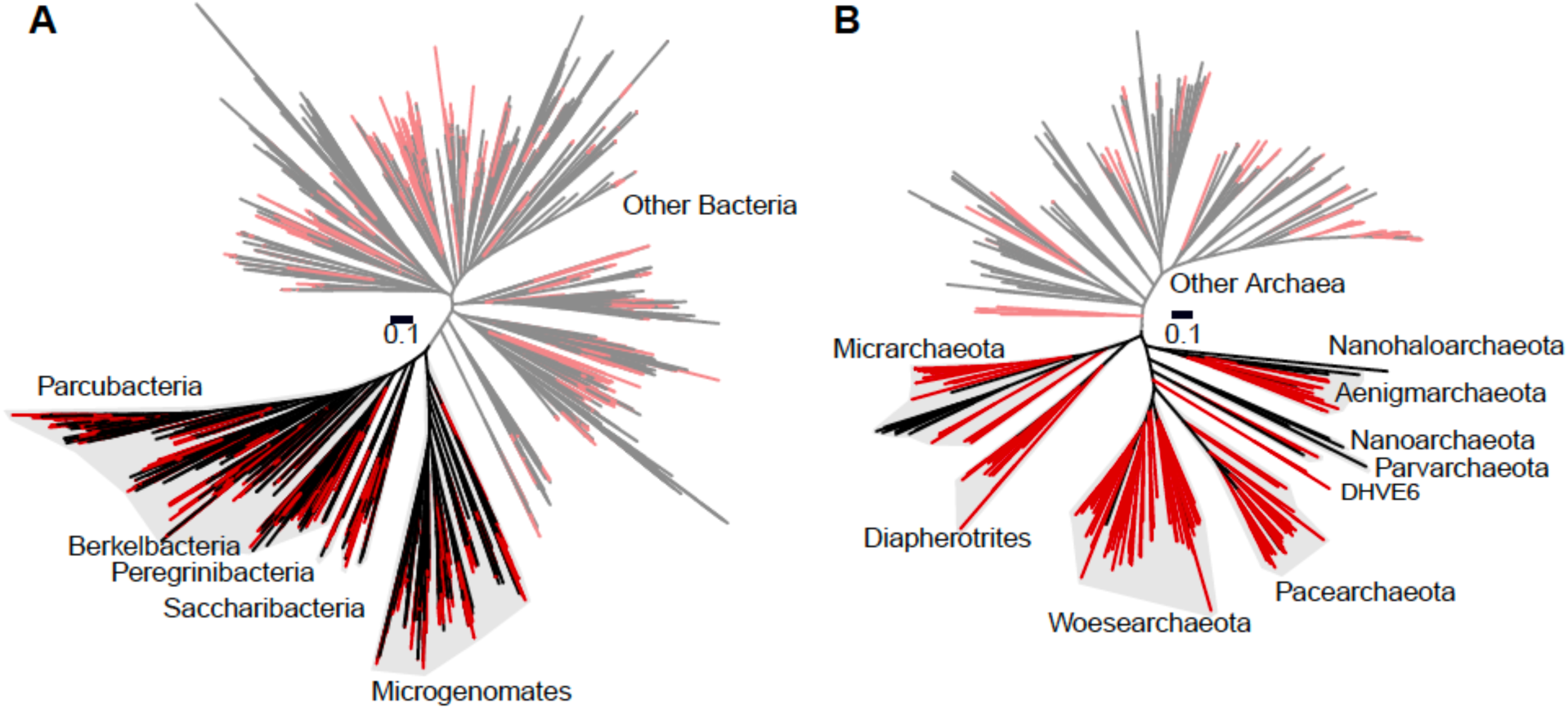
Maximum likelihood concatenated phylogenetic tree of single copy proteins (n=5) from Bacteria (a) and Archaea (b). All SAGs from this study are highlighted red. Patescibacteria and DPANN are highlighted with grey and labeled by individual proposed phyla within the superphylum (Rinke et al., 2013; Brown et al., 2015).

**Figure 3.**
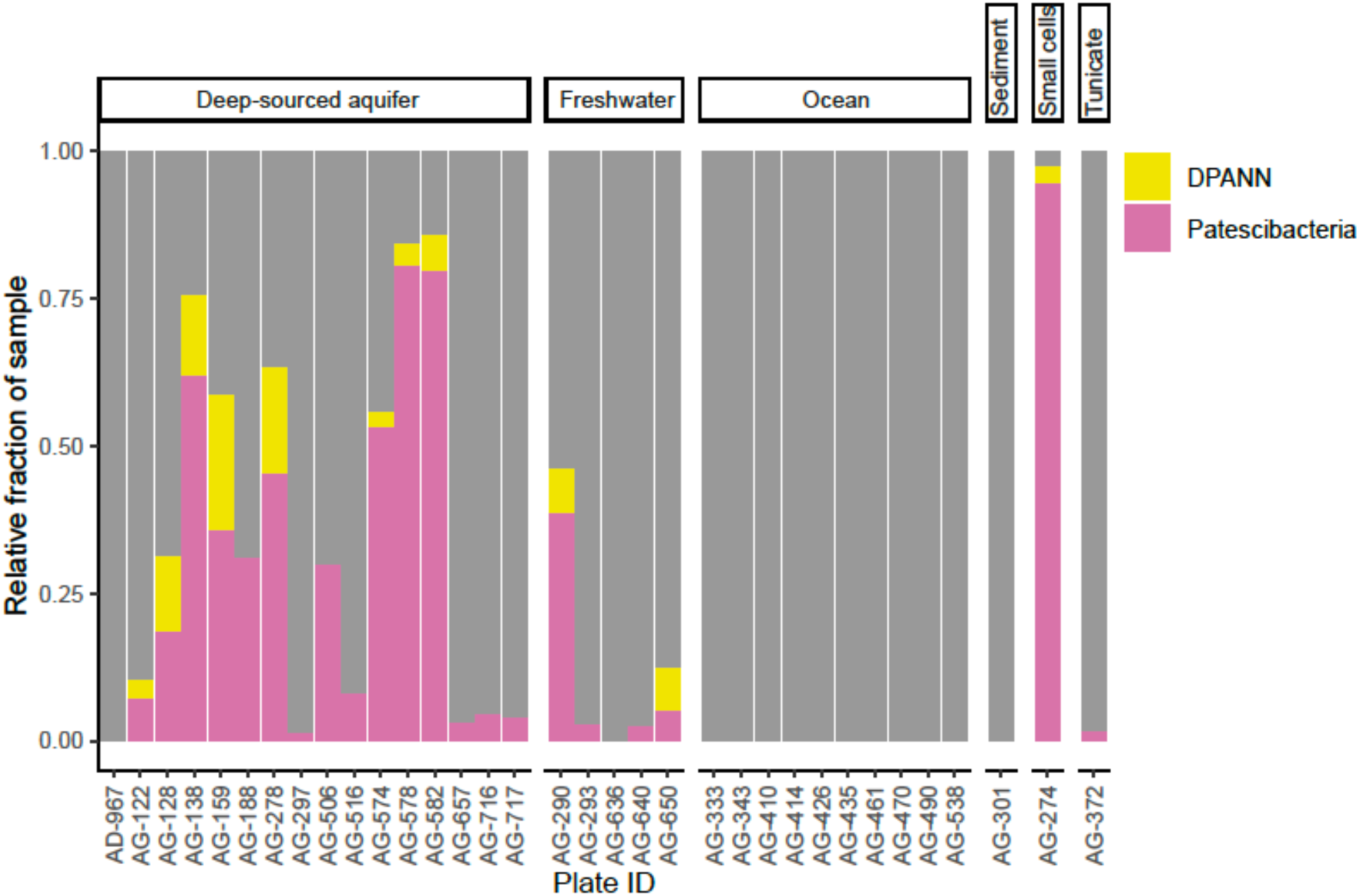
Relative abundance of Patescibacteria and DPANN from 34 geographically diverse samples determined from randomly sequenced LoCoS SAGs. The plate identifiers can be cross-referenced with specific SAGs and geographic sites in Table S1. The AG-274 sample contains small cells from a water-filled rock fracture at 1,340 m depth below surface in the Beatrix gold mine in South Africa.

### Evidence for physical cell-cell associations

We searched for evidence of physical cell-cell associations—an implication of obligate symbiosis—by identifying genomic sequences from multiple phylogenetically distinct organisms within individual SAGs. First, we searched for multiple copies of conserved, single copy protein-encoding genes using checkM (Parks et al., 2015), which is a commonly used tool to detect genome contamination. This approach identified 1% of Patescibacteria SAGs (5/492), 1.2% of DPANN SAGs (1/81), and 0.3% of SAGs from other phyla (5/1686) as containing DNA from heterogeneous sources (Table S3). Next, we searched for non-identical, near-full-length (> 1,000 bp) 16S rRNA genes in individual SAG assemblies. Such cases accounted for 1.5% of Patescibacteria (4/262), 0% DPANN (0/56), and 0.53% for other phyla (4/758) (Table S3). A Chi-square test revealed that there was a significant relationship between phyla and potential co-sorted SAGs from both checkM (p-value=1.2 x 10^−13^; Χ^2^=224.2) and 16S rRNA gene analyses (p-value<2.2 x 10^−16^; Χ^2^=238.07), but the overall contribution of Patescibacteria and DPANN to the significance of co-sorted SAGs was very low (<0.5%) relative to other phyla (Figure 4). Due to the incomplete SAG assemblies (Table S1), these sequencing-based approaches may underestimate the overall frequency of cell-cell associations in our data set. However, they consistently show that putative cell-cell associations constitute only a minor fraction of all SAGs, and that Patescibacteria and DPANN are not significantly enriched in such associations relative to other phyla in the studied environments. Furthermore, all identified cases of heterogeneous DNA in SAG assemblies were phylogenetically unique (Table S3), in contrast to the recurring Nanoarchaeota-Crenarchaeota symbiotic associations found using the same techniques in hot springs in prior studies (Munson-McGee et al., 2015; Jarett et al., 2018). Also, in mammalian oral microbiomes, Saccharibacteria have been shown to be specifically associated with Actinobacteria hosts (He et al., 2015; Cross et al., 2019). This suggests that the infrequent and inconsistent presence of taxonomically heterogeneous DNA in SAGs most likely originated from non-specific aggregation of multiple cells and/or attachment of extracellular DNA.

**Figure 4.**
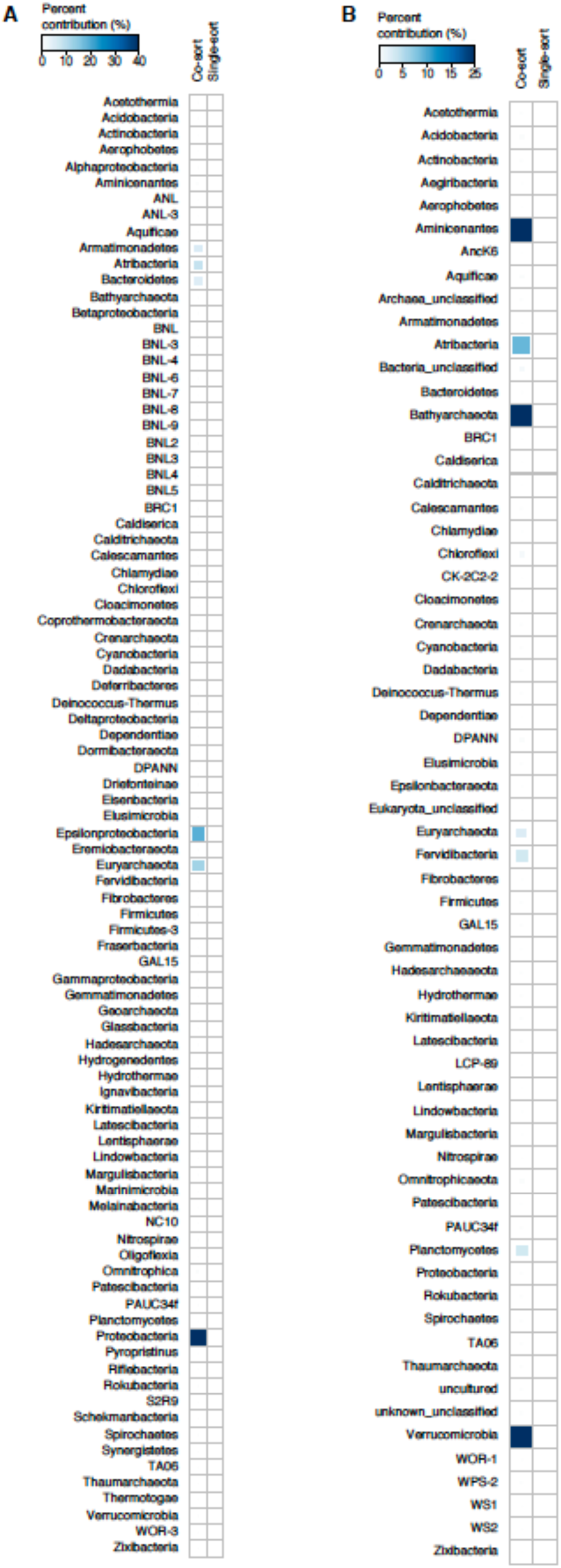
Plot of the percent contribution of individual phyla to the Chi-square statistic from checkM (**A**) and 16S rRNA gene (**B**) co-sorting analyses. Classification of phyla in (**A**) from concatenated phylogenetic tree in Figure 2 (Table S1) and from 16S taxonomy in (**B**).

Based on a small number of transmission electron micrograph observations, it has been suggested that Patescibacteria associations with other microorganisms may be fragile (Luef et al., 2015). Thus, we cannot rule out the possibility that some Patescibacteria and DPANN cells were attached to host cells *in situ* and became detached during sample collection and processing. To reduce the risk of dispersing natural cell aggregates and associations, we performed only a gentle mixing of the analyzed samples in preparation for cell sorting. In prior studies, similar techniques successfully revealed host-symbiont associations in termite guts (Hongoh et al., 2008), marine plankton (Martinez-Garcia et al., 2012) and hot springs (Jarett et al., 2018). This approach was also used to determine symbiotic associations between anaerobic methane-oxidizing archaea and their syntrophic partners in natural consortia from methane seeps (Hatzenpichler et al., 2016). It is worthy to note that the Saccharibacteria-Actinobacteria symbiont-host relationship was only disrupted by physical passage through a narrow-gauge needle multiple times (He et al., 2015). Also, putatively co-sorted SAGs of Nanoarchaeota and Crenarchaeota from iron oxide microbial mats were treated by a repeated physical disruption through multiple wash cycles and density gradient centrifugation, from which co-sorted cells were obtained (Jarett et al., 2018). Thus, although the techniques applied here may underestimate the overall counts of cell-cell associations *in situ*, we found no evidence for Patescibacteria and DPANN to be enriched in such associations relative to other phyla, and to form lineage-specific associations in the analyzed environments.

### Cell diameters

We employed calibrated index fluorescence-activated cell sorting (FACS) to determine physical diameters of individual cells that were used in SAG generation (Stepanauskas et al., 2017). This indicated that Patescibacteria (n=273) and DPANN (n=29) cells are extremely small across their entire phylogenetic breadth, with median estimated diameters of 0.2 µm (Figure 5). Several cases of larger, outlier diameter estimates may be due to attachment to other cells and particles, cellular division, methodological artifacts, or true biological variation. The low frequency of Patescibacteria and DPANN DNA recovery from larger particles (Table S1; Figure 5) provides further indication that most of these cells are not attached to other microorganisms. Likewise, most of the SAGs with identified heterogeneous genome sources were larger than their phylum median cell diameters (Table S3), which is consistent with their aggregation with other cells.

**Figure 5.**
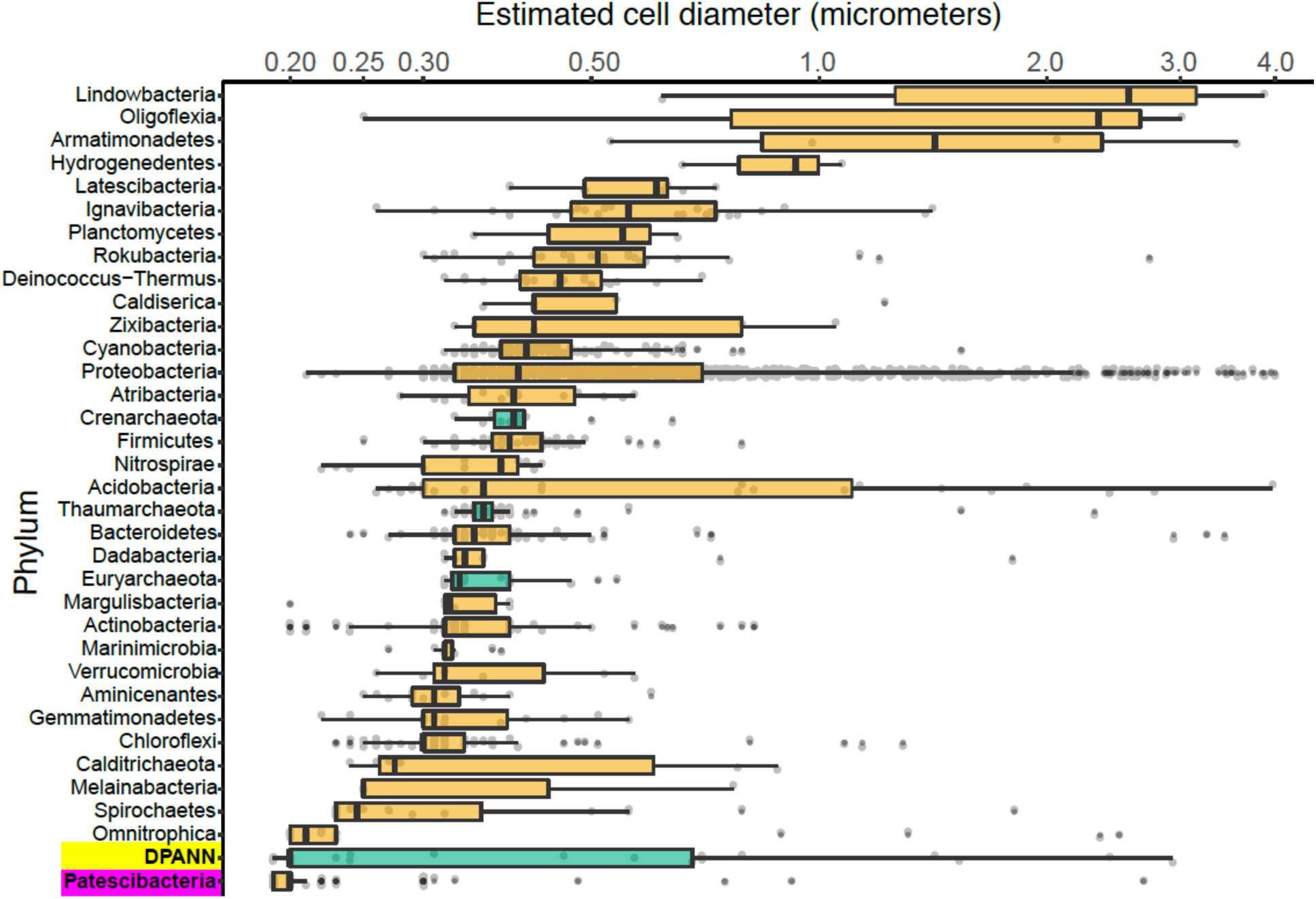
Phylum-resolved cell diameters. Solid black bars indicate medians; boxes represent the interquartile ranges (IQR) of the 1_st_ (Q1) and 3_rd_ (Q3) quartiles; whiskers denote the minimum (Q1 - 1.5*IQR) and maximum (Q3 + 1.5*IQR) values; outliers outside of the whiskers are marked by black dots. Orange indicates Bacteria and green indicates Archaea. A pairwise ranked-sum Wilcoxon test confirmed that the median diameter of Patescibacteria (highlighted in magenta) was smaller than most other phyla (27/36 phyla with p-values < 0.05; Table S5). The median diameter of DPANN (highlighted in yellow) was not significantly different from other archaea (1/36 phyla with p-values < 0.05; Table S5), likely due to the large variability in DPANN cell diameters. Individual cell diameters are available in Table S1 and pairwise p-values are located in Table S5.

To further investigate the composition of extremely small cells, we generated a complementary library of SAGs from a single subsurface sample (AG-274; Table S1) with a FACS gate targeting only ≤0.3 µm particles. Confirming our expectations, >90% of SAGs in this cell diameter-specific library were composed of Patescibacteria and DPANN (Figure 3). The obtained cellular size ranges are consistent with a prior report, which was based on transmission electron micrographs from one field study site (Luef et al., 2015). These cell diameters approximate the lower theoretical limits for cellular life (Maniloff et al., 1997).

### General genome features

To identify functional coding potential differences of Patescibacteria and DPANN compared to other Bacteria and Archaea, we performed a principal component analysis (PCA) using the relative abundance of clusters of orthologous groups (COG) as input variables with SAGs that had at least 30% completeness and a near full-length 16S rRNA gene (Figure 6). This showed a clear separation of Patescibacteria and DPANN from other bacteria and archaea along the first component (PC1) (Wilcoxon signed-rank test; p-value < 2.2 x 10^−16^). Importantly, well-described symbionts (Table S4) separated from both Patescibacteria along PC1 and DPANN along PC2 (p-value = 2.57 x 10^−8^ and 1.0 x 10^−7^ for Patescibacteria and DPANN, respectively). The only lineages that clustered with Patescibacteria and DPANN along PC1 and PC2 were Dependentiae and Tenericutes, respectively.

**Figure 6.**
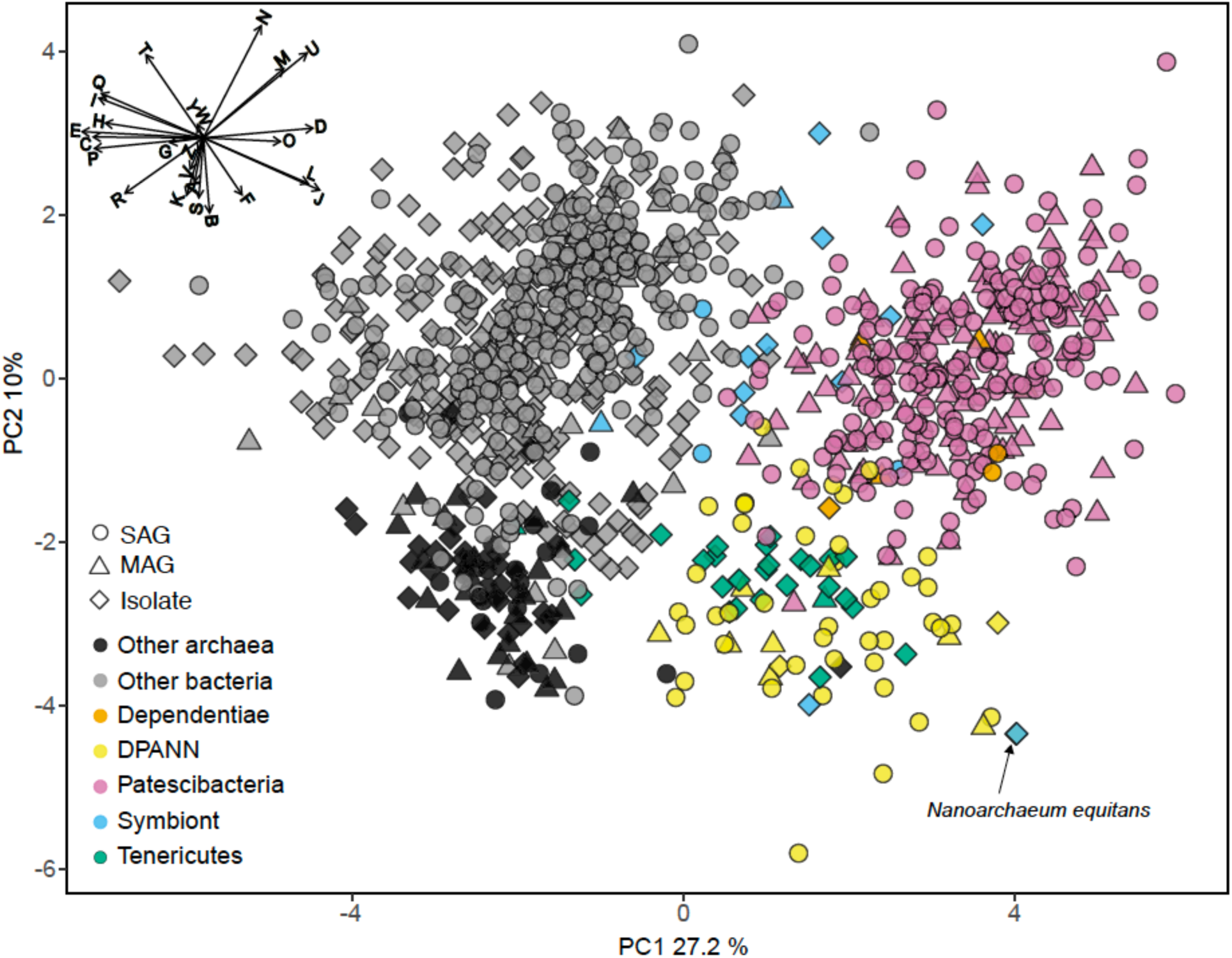
Principal components analysis (PCA) of the relative abundance of clusters of orthologous groups (COG) categories as the input variables. SAGs from this study (Table S1) and other studies (Table S2) with >30% completeness and had a near-full-length 16S rRNA gene and were included in the phylogenetic tree in Figure 8 (n=1,092). The vector plot in the upper left corner shows the COG categories that contributed to the most separation of the genomes: **Information Storage and Processing** Translation, ribosomal structure and biogenesis (J), RNA processing and modification (A), Transcription (K), Replication, recombination and repair (L), Chromatin structure and dynamics (B); **Cellular Processes and Signaling** Cell cycle control, cell division, chromosome partitioning (D), Nuclear structure (Y), Defense mechanisms (V), Signal transduction mechanisms (T), Cell wall/membrane/envelope biogenesis (M), Cell motility (N), Cytoskeleton (Z), Extracellular structures (W), Intracellular trafficking, secretion, and vesicular transport (U), Posttranslational modification, protein turnover, chaperones (O); **Metabolism** Energy production and conversion (C), Carbohydrate transport and metabolism (G), Amino acid transport and metabolism (E), Nucleotide transport and metabolism (F), Coenzyme transport and metabolism (H), Lipid transport and metabolism (I), Inorganic ion transport and metabolism (P), Secondary metabolites biosynthesis, transport and catabolism (Q),; **Poorly Characterized** General function prediction only (R), Function unknown (S). SAG, single amplified genome; MAG, metagenome assembled genome. Symbiont genomes are listed in Table S4. Note position of *Nanoarchaeum equitans* with black arrow.

The COG categories with the greatest negative effect on PC1, indicative of their relative depletion in Patescibacteria and DPANN, included E (amino acid metabolism and transport), C (energy production and conversion), P (inorganic ion transport and metabolism), and H (coenzyme transport and metabolism). The COG categories with the greatest positive effect on PC1, indicative of their relatively high fraction in genomes of Patescibacteria and DPANN, included D (cell cycle control and mitosis) and O (post-translational modification, protein turnover, chaperone functions). Archaea separated from bacteria along the second component (PC2) (p-value < 2.2 x 10^−16^) primarily by their relative enrichment in COG categories B (chromatin structure and dynamics), K (transcription), and S (unknown functions). This reflects the major inter-domain differences in DNA packing and transcription, and the greater fraction of archaeal genomes remaining uncharacterized, as compared to the genomes of Bacteria.

Genomes recently shaped by symbiosis often have low coding densities due to rapid gene loss and pseudogene formation (McCutcheon and Moran, 2012). Inconsistent with this pattern, we found the coding density of Patescibacteria and DPANN (median = 91%) to be typical of Bacteria and Archaea (median = 90%), while well-characterized symbionts were separated by their lower coding density (Figure 7a) (median = 0.87%, p-value = 0.035 and 0.028 compared to Patescibacteria and DPANN). Although the reduced genome size of Patescibacteria and DPANN has been viewed as an indication of a symbiotic lifestyle (Castelle et al., 2018), similar genome sizes (1-2 Mbp) are typical among free-living, marine plankton (Swan et al., 2013; Giovannoni et al., 2014). Furthermore, recent synthetic biology experimentation has pushed the minimal genome size limit of a free-living microorganism to ∼0.5 Mbp (Hutchison et al., 2016), far below the predicted sizes of Patescibacteria and DPANN genomes. Collectively, these general genome features of Patescibacteria and DPANN do not provide convincing evidence of an obligate symbiotic lifestyle.

**Figure 7.**
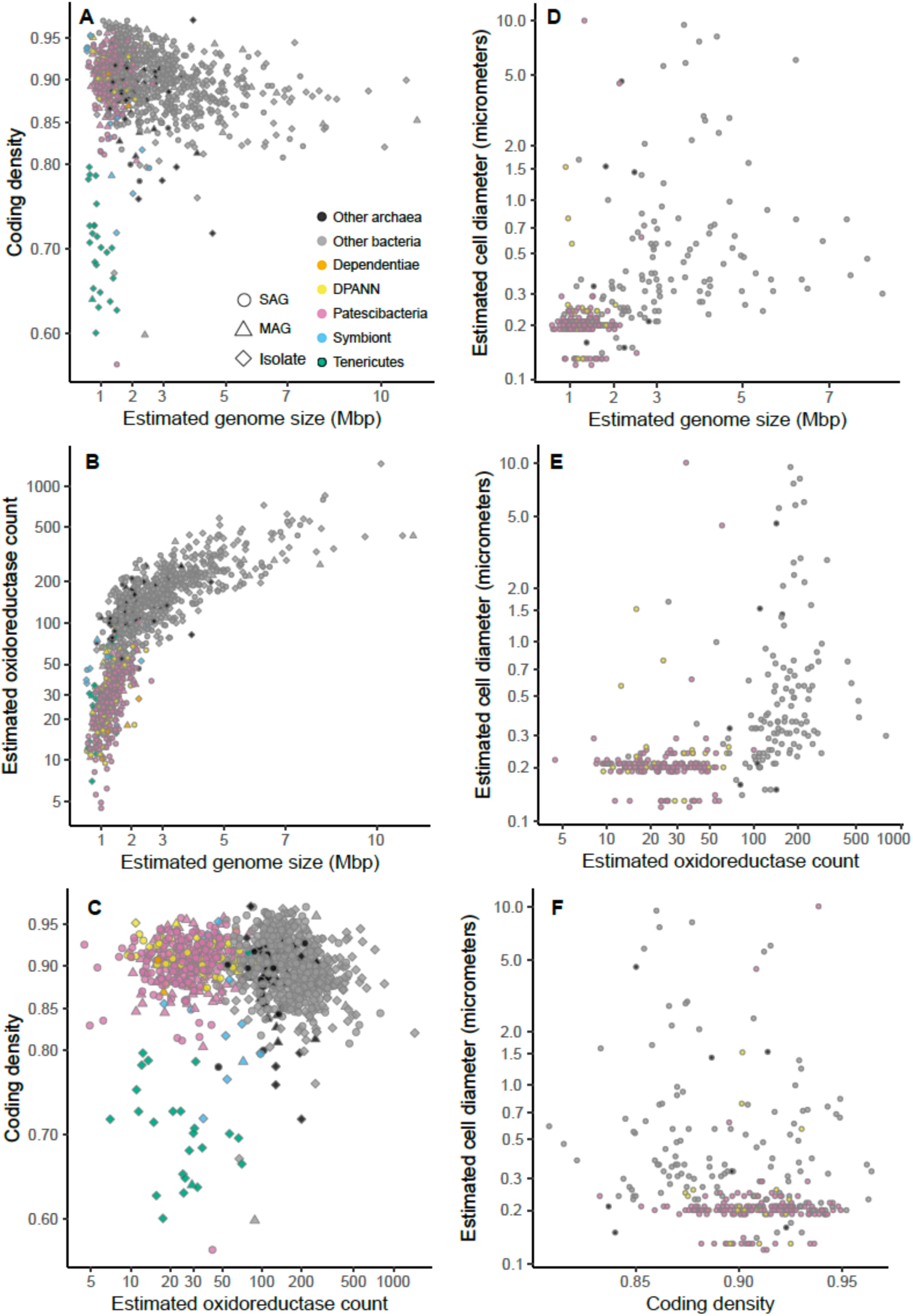
Relationship plots between estimated genome size, coding density, oxidoreductase count, and cell diameter among SAGs (Table S1) and other genome sequences (Table S2) that were greater than or equal to 30% complete, and were included in the 16S rRNA gene tree in Figure 8. Symbiont genomes are listed in Table S4.

In this context, the observed gene content similarities between Patescibacteria and Dependentiae, and between DPANN and most Tenericutes are intriguing (Figure 6). Dependentiae is a candidate bacterial phylum that has been noted for its reduced coding potential, including a depletion in electron transport chain components (McLean et al., 2013; Yeoh et al., 2016). It has been speculated that these characteristics indicate a symbiotic lifestyle, with energy acquired from hosts via ATP/ADP translocases, which has been confirmed experimentally in a few Dependentiae members (Delafont et al., 2015; Pagnier et al., 2015; Deeg et al., 2019). The well-characterized members of the bacterial phylum Tenericutes consist mostly of obligate pathogens with reduced genomes (Moran and Wernegreen, 2000). Interestingly, most Tenericutes are able to grow as free-living cells in rich media solely by fermentation (Tully et al., 1977), and were originally hypothesized to represent ancient lineages of life due to their small genome sizes and limited metabolisms (Morowitz, 1984). While we found all analyzed Dependentiae and most Tenericutes deplete in oxidoreductases (Figures 7b; Figure 8), only Tenericutes had a consistently low coding density (median = 71%) that is a characteristic of recently evolved symbionts (McCutcheon and Moran, 2012) (Figure 7a). Thus, we hypothesize that these two phyla cluster with the Patescibacteria and DPANN due to similar metabolic features arrived at by convergent evolutionary processes.

**Figure 8.**
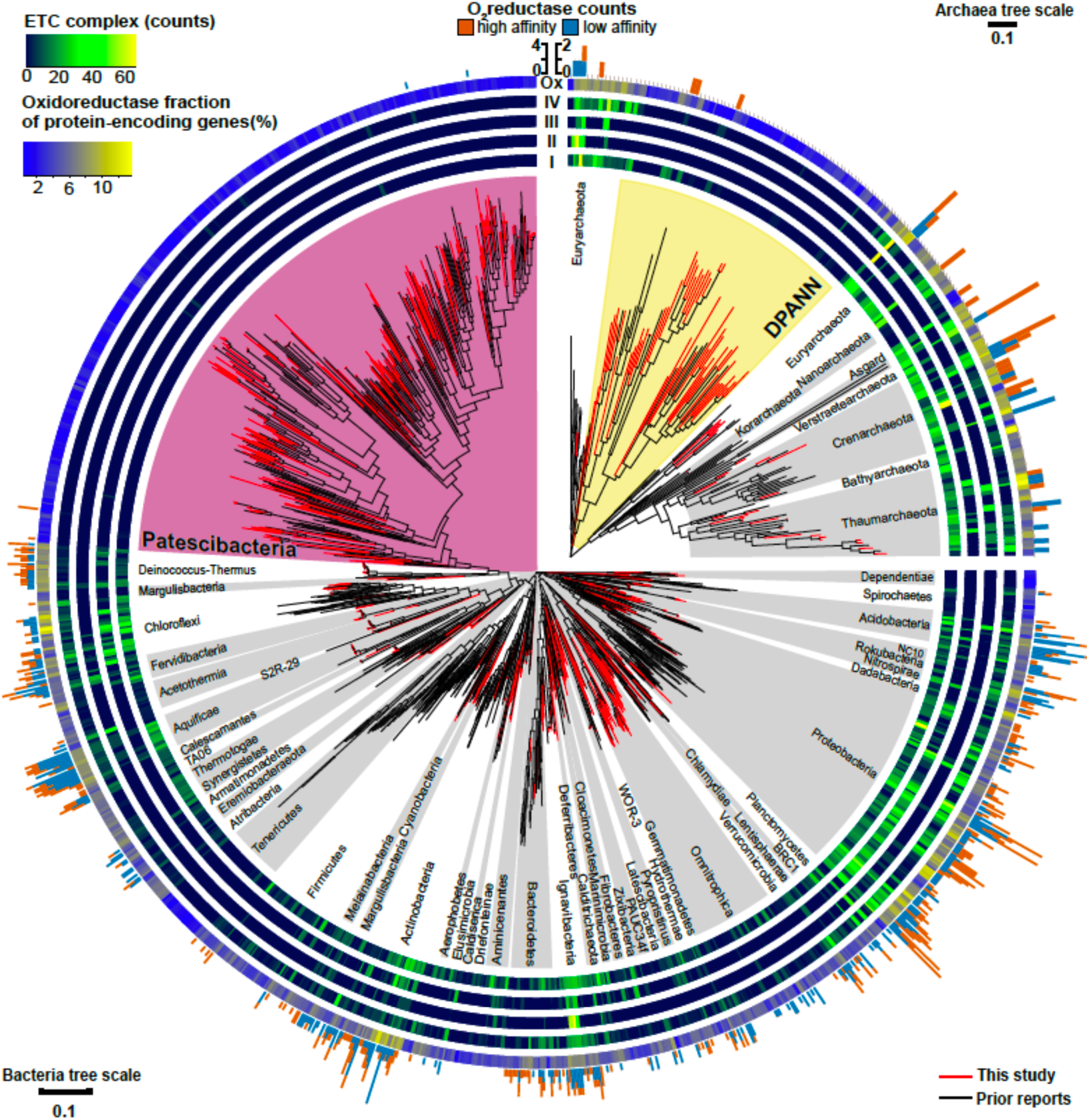
Maximum likelihood phylogeny of near full-length (>1,200 bp) 16S rRNA genes from Bacteria and Archaea, annotated with the distribution of counts for electron transport chain complexes, oxygen reductases, and oxidoreductase (Enzyme Commission 1; EC1) relative abundances from SAGs in this study (Table S1) and previously reported genome sequences (Table S2). The four innermost rings depict the counts of the electron transport chain complexes: I (NADH dehydrogenase subunits), II (succinate dehydrogenase subunits), III (cytochrome c reductase subunits), and IV (oxygen, nitrate, sulfate, iron, arsenate, and selenate reductase subunits). The outermost ring shows the relative abundance of oxidoreductases (Ox) for each genome assembly as a gradient from low (blue) to high (yellow). The peripheral stacked bar charts show the counts of oxygen reductases from both the heme copper oxidase and bd-ubiquinol oxidase oxygen reductase (O_2_red) families grouped as high (orange) or low (sky blue) affinity for oxygen (note scale bar differences between bacterial and archaeal trees). Patescibacteria are highlighted in magenta and DPANN are highlighted in yellow. Other bacterial and archaeal phyla are highlighted in alternating white and grey.

### Oxygen reductase genes

In search for an alternative explanation for the unique genealogy, genome content, and cell sizes of Patescibacteria and DPANN, we examined their energy metabolic coding potential. We found that only 0.6% of Patescibacteria SAGs (3/492) and none of the DPANN SAGs (0/81) from these samples encoded for homologs of oxygen reductases (O_2_red), as indicated by the presence of oxygen-binding subunit I of either the heme-copper oxidase (HCO) or bd-ubiquinol (bd) oxidase families. The incomplete genome recovery from individual SAGs cannot explain this pattern, because the 492 Patescibacteria SAGs and 81 DPANN SAGs correspond to a cumulative assembly of 162 and 27 randomly sampled, complete genomes. Furthermore, a phylogenetic analysis revealed that all three oxygen reductases from Patescibacteria SAGs form a cluster with other Patescibacteria sequences (Brown et al., 2015; Nelson and Stegen, 2015; León-Zayas et al., 2017) that is nested within a clade comprised of other phyla (Figure 9). We infer these phylogenetic relationships as an indication of a relatively recent horizontal gene transfer (HGT), likely from Proteobacteria and Firmicutes for the HCO and bd sequences, respectively. Although we did not detect any homologs of oxygen reductases in DPANN SAGs from our samples, the publicly available bd O_2_red sequences from DPANN metagenome bins and isolates formed a clade with Actinobacteria and Firmicutes, which we also infer as likely products of relatively recent HGT events. The topology of these O_2_red phylogenetic trees is consistent with prior reports, which have also been interpreted as evidence for prevalent HGT of oxygen reductase genes among other phyla (Brochier-Armanet et al., 2009; Gribaldo et al., 2009; Borisov et al., 2011). This suggests that the absence of oxygen reductases in Patescibacteria and DPANN is ancestral and not a result of gene loss due to adaptations to symbiosis, as previously hypothesized (Castelle et al., 2018).

**Figure 9.**
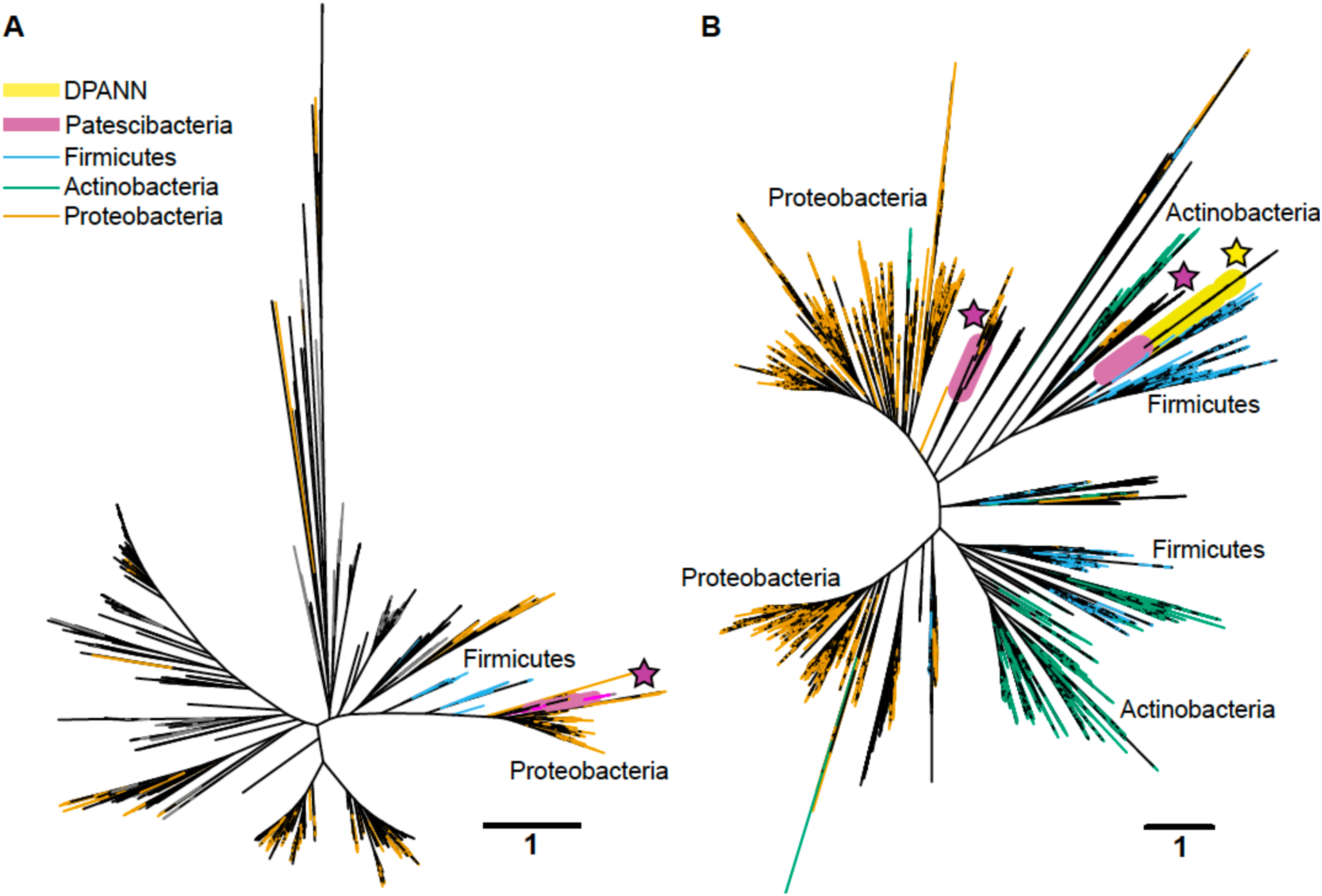
Maximum likelihood phylogenetic trees of the oxygen-binding subunit I from the heme copper oxidase (HCO) type A (a) and the A subunit from the bd-ubiquinol (b) oxygen reductase families. Patescibacteria and DPANN sequences are marked with magenta and yellow stars, respectively. The Patescibacteria HCO type A sequences (a) are nested within a larger clade containing mostly Proteobacteria (orange), and the Patescibacteria and DPANN bd-ubiquinol sequences (b) are nested within Proteobacteria (orange) and Firmicutes (blue) dominated clades. The scale bar represents the estimated number of substitutions per site

### Distribution of electron transport chain complexes

Patescibacteria and DPANN were depleted in the entire family of oxidoreductase enzyme genes compared to other bacteria and archaea, (p-value < 2.2 x 10^−16^) (Figures 7b, 8). This depletion was also significant in relation to symbionts with their comparatively small genome sizes (p-value < 0.05). Oxidoreductases are key components of both aerobic and anaerobic respiratory pathways (Jelen et al., 2016), so underrepresentation of them would suggest reduced functionality of these energy transduction mechanisms. Accordingly, none of the Patescibacteria and DPANN genomes were found to encode a complete ETC consisting of all four complexes (Figure 8). Putative homologs of at least two of the four ETC complexes were found only in 3% and 11% of Patescibacteria and DPANN genomes, respectively. We found putative homologs of genes encoding individual complexes I, II, III, and IV in 0%, 2%, 3%, and 14% of Patescibacteria genomes. The corresponding numbers for DPANN were 7%, 4%, 0%, and 21%. Some of these computationally predicted genes are only distantly related to experimentally verified homologs and therefore may constitute false positives. These findings are consistent with the lack of complete ETC reports in prior studies of Patescibacteria genomes (Brown et al., 2015), with the sole exception of a tentative nitric oxide respiration operon found in a single metagenome bin (Castelle et al., 2017). The sparse and scattered distribution of the putative ETC gene homologs in Patescibacteria and DPANN (Figure 8) suggest horizontal gene transfer origins rather than ancestral inheritance. This is consistent with the phylogenetic reconstructions of other energy transducing genes identified in Patescibacteria, which also suggest evolutionary origins from horizontal gene transfer (Jaffe et al., 2019). Collectively, our observations indicate that the absence of complete electron transport chains in Patescibacteria and DPANN is an ancestral feature, which we propose is more parsimonious than multiple gene loss events due to obligate symbiosis (Brown et al., 2015; Hug et al., 2016; Castelle et al., 2018; Dombrowski et al., 2019; Méheust et al., 2019).

### Respiration activity

To experimentally test for the presence of active oxidoreductases in a subsurface microbial community, we employed the fluorogenic oxidoreductase probe RedoxSensor Green on a deep groundwater sample from South Dakota. This revealed a wide range in fluorescence intensity in phylogenetically diverse cells, with none of the Patescibacteria cells exceeding the fluorescence of particles in a heat-killed, negative control (Figure 10). To the best of our knowledge, RedoxSensor Green has not been tested extensively on diverse microbial lineages, therefore these results should be considered tentative. Nonetheless, both genome content and *in situ* physiology analyses indicate the absence of respiration in Patescibacteria and DPANN, which corroborates earlier reports of these lineages containing few, if any, components of energy transducing pathways other than fermentation (Castelle et al., 2018).

**Figure 10.**
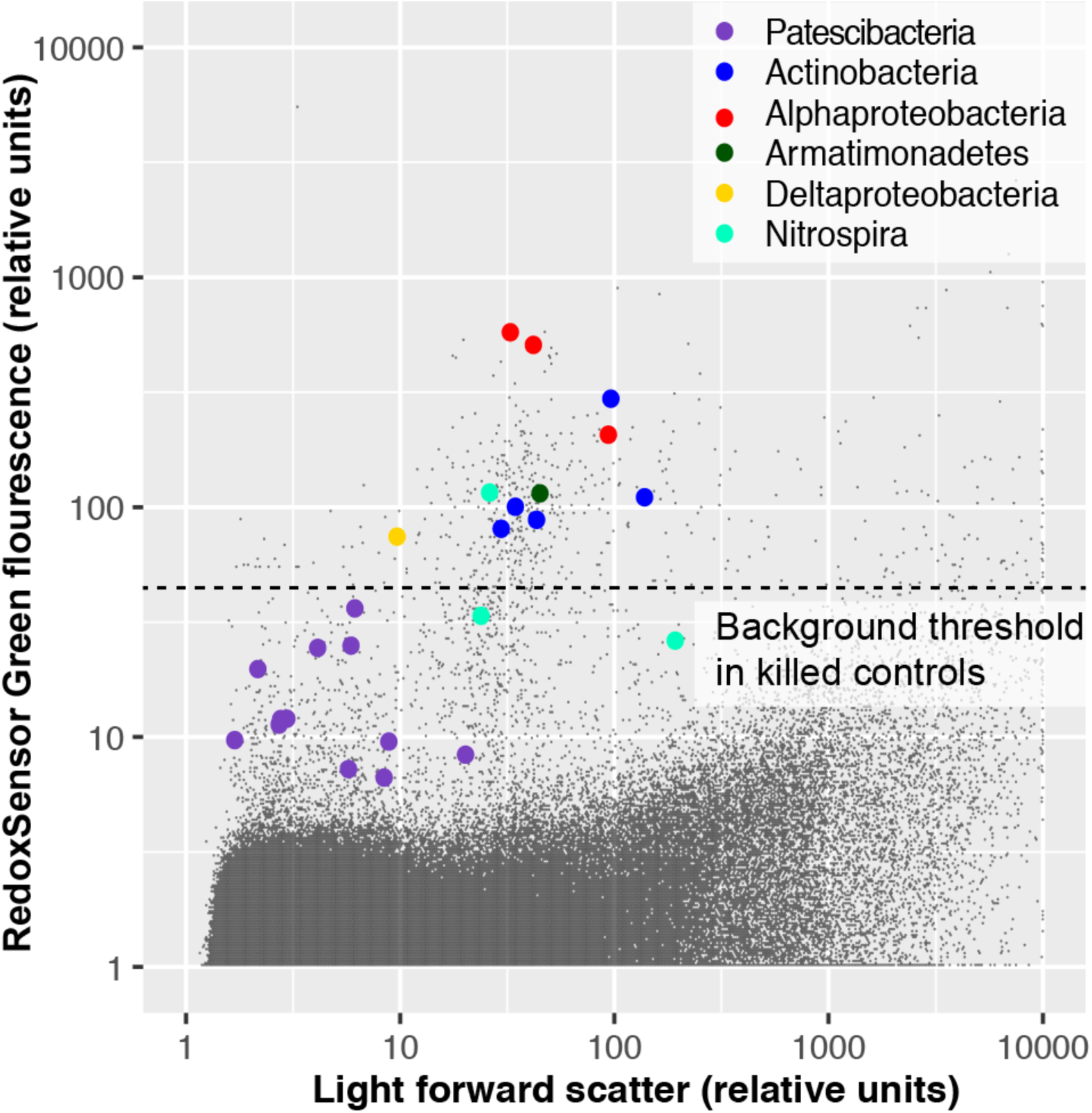
Oxidoreductase activity in subsurface (∼300 m below surface) microbial cells from Homestake Mine (Lead, South Dakota, USA) measured by RedoxSensor Green (RSG; ThermoFisher).

### 16S rRNA gene phylogeny

The placement of Patescibacteria and DPANN in the tree of life is widely debated (Hug et al., 2016; Williams et al., 2017; Dombrowski et al., 2019). Most current phylogenetic inferences are based on concatenated single-copy proteins (CSCP), which has the advantage of higher phylogenetic resolution, as compared to phylogenies of individual genes (Rinke et al., 2013). However, the unknown genetic change at heterogeneously evolving sites and large sequence divergence may limit the accuracy of such trees (Pace, 2009; Dombrowski et al., 2019). To complement the CSCP-resolved genealogy (Figure 2), we performed a large-scale phylogenetic analysis of the well-established 16S rRNA gene (Woese, 2002) (length > 1,200 bp) separately for Bacteria and Archaea. The obtained phylogenetic inference (Figure 8) supported the separation of Patescibacteria and DPANN from other bacterial and archaeal lineages, in agreement with the phylogenies based on CSCP genes (Castelle et al., 2018) (Figure 2) and a recent large scale bacterial 16S rRNA gene tree (Schulz et al., 2017). Importantly, we did not observe grouping of Patescibacteria with fast-evolving lineages (e.g., obligate insect symbionts and Tenericutes) that could be due to long branch attraction in the 16S rRNA gene phylogeny. This suggests that the divergent branching of Patescibacteria and DPANN is probably not a result of recent, accelerated divergence.

## Concluding Remarks

Using the collective evidence from cell-cell association, coding potential and phylogenomic analyses, we propose a new explanation of the unusual biological features of Patescibacteria and DPANN. Although the Patescibacteria and DPANN contain symbionts (Huber et al., 2002; Podar et al., 2013; Gong et al., 2014; He et al., 2015; Munson-McGee et al., 2015; Jarett et al., 2018; Cross et al., 2019; Hamm et al., 2019) and auxotrophies (Castelle et al., 2018; Dombrowski et al., 2019), we believe that there is not sufficient evidence to conclude that an ancestral adaptation to symbiosis has led to the reduction of their cell sizes and coding potential (Castelle et al., 2018; Dombrowski et al., 2019; Méheust et al., 2019). Instead, our data indicate that most Patescibacteria and DPANN do not form symbiotic cell-cell associations in subsurface environments, and that their divergent coding potential, small genomes, and small cell sizes may be a result of a primitive energy metabolism that relies solely on substrate-level phosphorylation (fermentation), potentially preceding the evolution of electron transport phosphorylation (respiration). Auxotrophies are very common among microorganisms, and represent a wide range of dependencies for exogenous cellular components (Zengler and Zaramela, 2018). Patescibacteria and DPANN may be on the extreme end of the spectrum in their dependence on other community members, perhaps a reflection of an ancient evolutionary strategy to limit cellular biosynthetic energy requirements, as energetic allocation is a major driver of genome evolution in bacteria and archaea (Lynch and Marinov, 2015).

The authors declare no conflict of interest.

## Acknowledgements

We thank the staff of the Bigelow Laboratory for Ocean Science Single Cell Genomics Center (SCGC) for the generation of single cell genomic data. We also thank Paul Falkowski and Saroj Poudel for advice on electron transport systems, Jacob Munson-McGee for insightful discussions on co-sorted SAGs, and David Emerson, Sean Crowe, Paul van der Wielen, Sari Peura, Andreas Teske, Takuro Nunoura, Christa Schleper, and Steffen Joergensen for contributing field samples. Thanks also to Richard Friese, Josh Hoines, and Kevin Wilson (National Park Service); Michael King and John Bredehoft (Hydrodynamics Group); Darrell Lacy, Levi Kryder, John Klenke, and Jamie Walker (Nuclear Waste Repository Project Office); and Jaret Heise, Tom Regan and others at Sanford Underground Research Facility (SURF); Corey Lee (US Fish and Wildlife Service) and The Nature Conservancy for site access and sampling assistance in Nevada and South Dakota. Special thanks to Brittany Kruger, Josh Sackett, Scott Hamilton-Brehm, John Healey, Brad Lyles, and Chuck Russell of DRI for sampling assistance in Nevada. Thanks to the U.S. Forest Service for a permit to conduct research at Little Hot Creek, California and the 2015 International Geobiology Course for field work and sample collection. We also thank Vitaly Kadnikov, Olga Karnachuk and Tamara Zemskaya for sample collection in Siberia. Also, thanks to Stephen Grasby, Allyson Brady, and Christine Sharp for sampling Canadian field samples, and Wen-Jun Li for logistical support and access to hot spring samples in China. We also thank British Columbia Parks and Parks Canada for permission to sample Dewar Creek and Paint Pots field sites. This work was funded by the USA National Science Foundation grants 1441717, 1826734, and 1335810 (to RS); and 1460861 (REU site at Bigelow Laboratory for Ocean Sciences). TW, FS and JJ were funded by the U.S. Department of Energy Joint Genome Institute, a DOE Office of Science User Facility supported under Contract No. DE-AC02-05CH11231. NVR group was funded by the Russian Science Foundation (grant 19-14-00245). SMS was funded by USA National Science Foundation grants OCE-0452333 and OCE-1136727. BPH was funded by NASA Exobiology grant 80NSSC17K0548.

## Author contributions

JPB led data analyses and manuscript preparation. RS developed the concept and managed the project, with contributions by TW, TCO, DM, JAE, JPB and EDB. EDB, JMB, FS, JKJ, OB, KC contributed to data analyses. NJP performed cell sorting and size calibration at Bigelow Laboratory. TCO, DPM, PD, NVR, JRS, BPH, KAK, SMS, MSE, HAB and MBS oversaw field sample collection. All authors contributed to data interpretation and manuscript preparation.

## Supplemental Table Captions

**Table S1.** Deep-sequenced and LoCoS SAGs from this study with genomic statistics and associated environmental metadata. Data are ordered with the following column headers:

1=genome; single amplified genome (SAG identifier)

2=gold.analysis.id; Gold analysis identifier (used to search genome in IMG/M)

3=phylum

4=assembly.completeness; checkM completeness estimates

5= contamination; checkM estimated genome contamination

6=assembly.size; SAG assembly size

7=est.genome.size; estimated genome size

8=coding.density

9=ec1.count; counts of oxidoreductases from SAG assembly

10=est.ec1.count; estimated counts of oxidoreductases from predicted genome size of n Mbp

11=16s.copy.number; number of predicted 16S rRNA genes

12=cell.diameter; estimated cell diameter

13=sequencing.center

14=sample.collection.site; name of site where samples were collected

15=sample.type

16=date.collected; sample collection date dd/mm/yy

17=latitude

18=longitude

19=depth

20=dissolved.oxygen (micromoles/L)

21=ph

22=salinity (practical salinity units, psu)

23=temperature (degrees Celsius

24=h2s; dissolved hydrogen sulfide (millimoles/L)

NA=not applicable

**Table S2.** Genomes from other studies with associated genomic statistical information (accessed from IMG/M on April 2018).

**Table S3.** Potential co-sorted SAGs from deep-sequenced and LoCoS datasets. SAGs can be cross-referenced for specific information with Table S1.

**Table S4.** Symbiont genome assemblies and taxonomic names used in Figures 6 and 7.

**Table S5.** Pairwise Wilcoxon’s test p-values on all phyla versus phyla cell diameter estimations in Figure 5.

