## Supplemental Table 2 for "Ancestral absence of electron transport chains in Patescibacteria and DPANN"

| SAG | Method | Sequencing | Phylum association | Sample | Estimated cell diameter (µm) |
|---|---|---|---|---|---|
| JGI_BlankSpring-1-M16 | 16S | Deep | Patescibacteria-Euryarchaeota | Blank_Spring_Wilbur_Springs | nd |
| JGI_DC4.SYBR.4.E13.16S | 16S | Deep | Fervidibacteria-Fervidibacteria | Dewar_Creek | nd |
| JGI_Kivu325.minimeta.G3 | 16S | Deep | Patescibacteria-Atribacteria | Lake_Kivu | nd |
| JGI_Kivu325.minimeta.H3 | 16S | Deep | Atribacteria-Atribacteria | Lake_Kivu | nd |
| SCGC_AG-128-I19 | 16S | Deep | Patescibacteria-Planctomycetes | Ash_Meadows_Crystal_Spring | nd |
| SCGC_AG-128-I19 | 16S | LoCoS | Patescibacteria-Planctomycetes | Ash_Meadows_Crystal_Spring | nd |
| SCGC_AG-372-K08 | 16S | LoCoS | Proteobacteria-Bacteroidetes | Tunicate | 3.43 |
| SCGC_AG-470-F22 | 16S | LoCoS | Proteobacteria-Cyanobacteria | Ocean | 1.54 |
| SCGC_AG-582-A14 | 16S | Deep | Patescibacteria-Patescibacteria | SURF_1700_D_ferric_finger | 0.2 |
| SCGC_AG-650-J16 | 16S | Deep | Proteobacteria-Ignavibacteria-Proteobacteria | Little_Hot_Creek_Cone_Pool | 2.37 |
| SCGC_AH-147-O09 | 16S | Deep | Acetothermia-Verrucomicrobia | BLM | nd |
| JGI_DC4.SYBR.3.C22.16S | checkM | Deep | Aquificae-Aquificae | Dewar_Creek | nd |
| JGI_DC4.SYBR.3.G20.16S | checkM | Deep | Fervidibacteria-unknown | Dewar_Creek | nd |
| JGI_Kivu325.minimeta.G3 | checkM | Deep | Patescibacteria-Atribacteria | Lake_Kivu | nd |
| JGI_Kivu325.minimeta.H3 | checkM | Deep | Atribacteria-Atribacteria | Lake_Kivu | nd |
| JGI_Kivu325.minimeta.J3 | checkM | Deep | Aminicenantes-Chloroflexi | Lake_Kivu | nd |
| SCGC_AD-726-L19 | checkM | Deep | Patescibacteria-Epsilonproteobacteria | Crab_Spa | nd |
| SCGC_AG-372-K21 | checkM | Deep | Patescibacteria-Bacteroidetes | Tunicate | 3.25 |
| SCGC_AG-640-D13 | checkM | Deep | Patescibacteria-Aminicenantes | Zodletone_Spring | 0.27 |
| SCGC_AG-650-J04 | checkM | Deep | Patescibacteria-Armatimonadetes | Little_Hot_Creek_Cone_Pool | 4.48 |
| SCGC_AG-650-J16 | checkM | Deep | Ignavibacteria-Proteobacteria | Little_Hot_Creek_Cone_Pool | 2.37 |
| SCGC_AG-650-P07 | checkM | Deep | DPANN-Euryarchaeota | Little_Hot_Creek_Cone_Pool | 36.68 |

nd=no data
