## Supplemental Table 4 for "Ancestral absence of electron transport chains in Patescibacteria and DPANN"

| IMG Genome identifier | Phylum | Taxonomic name |
|---|---|---|
| 2512564067 | Proteobacteria | Rickettsia montanensis |
| 2568526683 | Proteobacteria | Holospora obtusa |
| 2576861681 | Proteobacteria | Candidatus Xenolissoclinum pacificiensis |
| 2609460328 | Proteobacteria | Candidatus Hepatobacter penaei |
| 2616645016 | Proteobacteria | Candidatus Endoecteinascidia frumentensis |
| 2630968813 | Bacteroidetes | Candidatus Sulcia muelleri |
| 2636415932 | Bacteroidetes | Cardinium endosymbiont cEper1 |
| 2740892203 | Spirochaetes | Treponema endosymbiont D11 |
| 637000036 | Spirochaetes | Borrelia burgdorferi |
| 637000331 | Actinobacteria | Tropheryma whipplei |
| 642555114 | Bacteroidetes | Candidatus Amoebophilus asiaticus |
| 642555127 | Elusimicrobia | Elusimicrobium minutum |
| 642555145 | Proteobacteria | Orientia tsutsugamushi |
| 646564518 | Bacteroidetes | Sulcia muelleri |
| 648028014 | Bacteroidetes | Sulcia muelleri |
| 650716011 | Bacteroidetes | Blattabacterium sp. Bge |
| 638154511 | DPANN | Nanoarchaeum equitans |
